# In silico design of bioactive chimeric peptide from archaeal antimicrobial peptides

**DOI:** 10.1101/2021.08.14.456327

**Authors:** Souvik Banerjee, Soham Chakraborty, Kaustav Majumder

**Affiliations:** Department of Microbiology, St. Xavier’s College (Autonomous), Kolkata, West Bengal, India; Department of Chemistry, Indian Institute of Technology, Indian School of Mines (ISM), Dhanbad, Jharkhand, India; Department of Biosciences and Bioengineering, Indian Institute of Technology, Bombay, Mumbai, Maharashtra, India

**Keywords:** Peptide therapeutics, multi-drug resistant bacteria, chronic human diseases, archaeal, computational tools

## Abstract

Novel peptide therapeutics have been the cardinal part of modern-day research. Such therapies are being incorporated to prevent the adverse effects of globally emerging multi-drug resistant bacteria and various chronic human diseases which pose a great risk to the present world. In this study, we have designed a novel peptide therapy involving archaeal antimicrobial peptides. In silico predictions assign the peptide construct to be antigenic, non-allergenic, non-toxic and having stable physicochemical properties. The secondary and tertiary structures of the construct were predicted. The tertiary structure was refined for improving the quality of the predicted model. Computational tools predicted intracellular receptors in *Escherichia coli*, *Klebsiella pneumoniae* and the human body to be possible binding targets of the construct. In silico docking of modelled peptide with predicted targets, showed prominent results against targets for complex human diseases and that of bacterial infections. The stability of those docked complexes was confirmed with computational studies of conformational dynamics. Certainly, the designed peptide could be a potent therapeutic against multi-drug resistant bacteria as well as several human diseases.

## Introduction

The first conventional peptide therapy dates back to insulin as a therapeutic for diabetes in the 1920s. Since then, the usage of peptides as drugs has morphed and evolved with time being continued to develop along the alterations in the modern day’s therapeutic paradigms. Usage of peptides as cost effective drugs had been thwarted for a long time due to their short plasma life, challenges in oral administration, and hydrophilic nature (physiological obstacles). Despite such restraints, experimental evidences have asserted their emergence as potential alternatives showing greater efficacy in selectivity and specificity in comparison to small molecule inhibitors (Muttenthaler et al. 2021). Peptide drugs, unlike their synthetic counterparts, degrade into corresponding amino acids inside the body, eschewing the formation of toxic metabolites. In addition to that, peptide drugs having short half-lives lead to less accumulation of degraded products in the body. (Hoppenz et al. 2020; Timmur and Gürsoy 2021).

The naturally occurring antimicrobial peptides (AMPs) having broad spectrum of action like cationic Polymyxin B, Gramicidin S and the cationic Lantibiotic nisin, are currently being used as efficient therapeutics to prevent the detrimental effects of antimicrobial drug resistance. AMPs are short length (ribosomally or non-ribosomally synthesized), structurally diverse, cationic peptides, present in large no. of organisms ranging from microbes to human. The list of existing effective AMPs is accessible through CAMPR3 database (Vale et al. 2014; Waghu et al. 2016). As a matter of fact, biological threats augmented by the increasing resistance to existing antimicrobial drugs has brought the usage of AMPs into light. AMPs are the need of the hour since the drug resistant bacterial infections and associated complexities have turned out to be a serious global health concern (Haney et al. 2018). The biofilm formations play a crucial role in increasing their resistance to antibiotic therapies and ability to evade host immune response by assisting the microbes to delve into persistent colonization in new niche and diverse environments (Majumdar and Pal 2017). In addition to that, biofilms ensuring survival in hostile environments has paved the way to the development of chronic infections leading to cancer predominantly gastric, pancreatic and mucinous colorectal carcinogenesis (Rizzato et al. 2019; Li et al. 2019). The most gripping standpoint of AMPs in such cases has been evinced experimentally in their ability to participate in host defence mechanism to colon carcinogenesis, bringing balance to the colon microbiome (Zhang et al. 2019).

Concomitant to the AMPs are the bacteriocins which are peptide toxins, produced by a bacterium to inhibit the spread of other bacteria. Hence bacteriocins show antimicrobial properties (Mathur et al. 2017). The most riveting features are of the archaeosins, the group of AMPs produced by Archaea (a diverse and abundant family of prokaryotes, also termed as ‘extremophiles’ due to their drastic living habitats). Archaeosins enable archaeabacteria to colonize in harsh conditions inhibiting the growth of other bacteria. The archaeosins are the sole ribosomally synthesized positively charged AMPs (Mahlapuu et al. 2016; Candido et al. 2019) to be experimentally characterized and identified only in some species of *Halobacteria* and *Sulfolobales* (Makarova et al. 2019). Taking into account this facet of archaeal biology, this research work unfolds a new dimension of preparing a potent curative with archaeal antimicrobial peptides which has not been studied before. Two peptide chains with prominent antimicrobial property were chosen from the archaeabacteria *Methanosarcina acetovorans* C2A, and *Sulfolobus acidocaldarius* DSM 639 as provided by the CAMPR3 database. Using these two archaeal AMPs, this scientific endeavour focuses scrupulously on the designing of a novel peptide therapeutic construct against bacterial infections as well as human diseases.

## Materials and methods

### Prediction of antimicrobial motifs in selected AMPs

The CAMPR3 database (http://www.camp.bicnirrh.res.in/seqDb.php) was accessed to obtain two archaeal AMPs with peptide length in the range 0 – 200 amino acids. Among the obtained peptide sequences, one was from *Methanosarcina acetovorans* C2A and the other from *Sulfolobus acidocaldarius* DSM 639. The acquired sequences were given as input in the antimicrobial region prediction (ARP) tool (http://www.camp.bicnirrh.res.in/predict_c/) of the database to analyze the major antimicrobial motifs. The peptide length parameter in ARP tool was set to the default value of 20 and the AMP probability was chosen as result sorting option. The SVM algorithm was selected for prediction. Both the predicted sequences consisting of common Linocin M18 (bacteriocin) motif were linked with a GGSG linker to form a recombinant construct (Fig. 1).

**Fig. 1.**
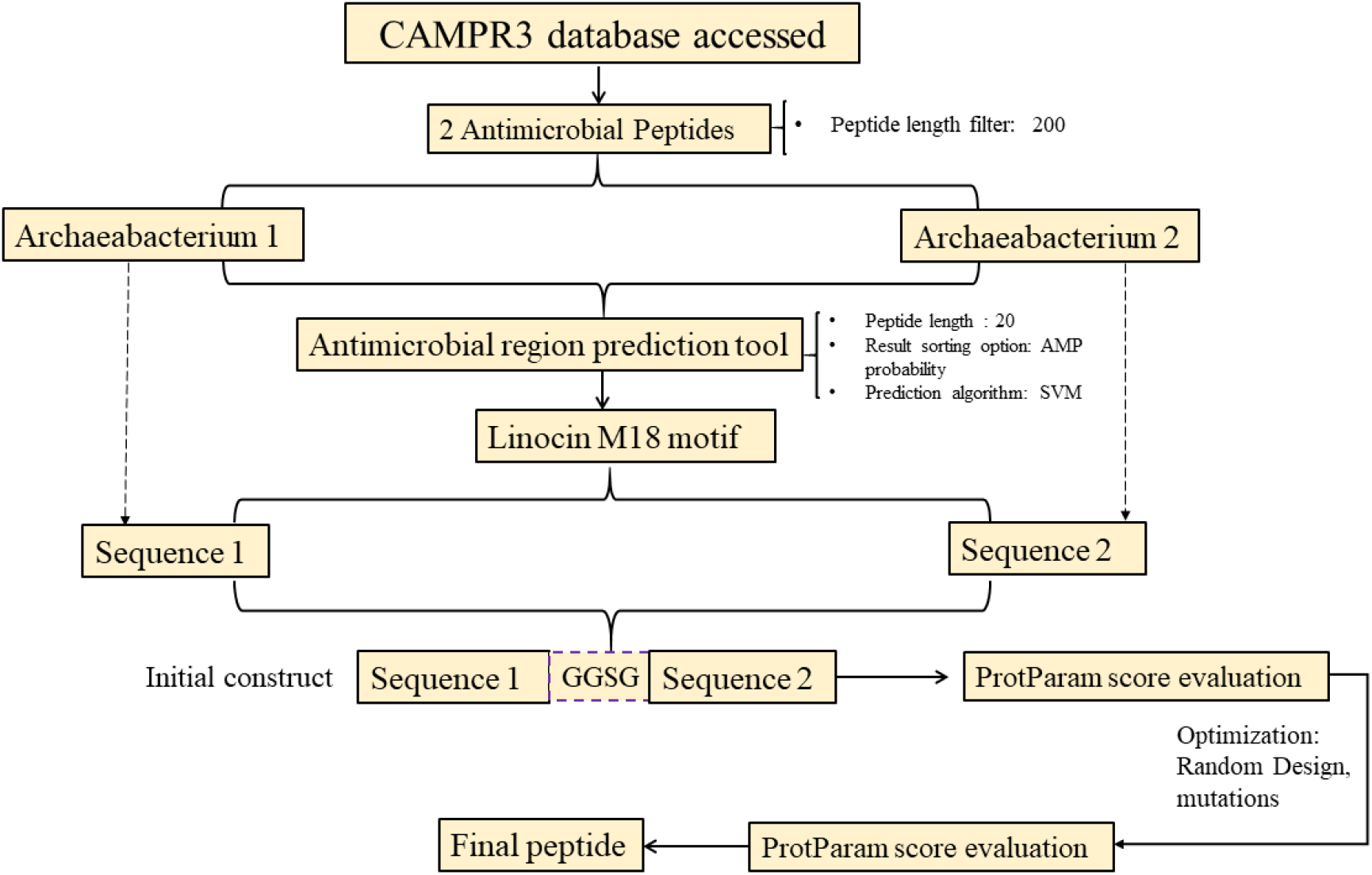
Workflow for designing the peptide construct

### Study of physicochemical properties of recombinant construct

The physicochemical properties of the prepared construct were analysed using the ProtParam tool (https://web.expasy.org/protparam/). The ProtParam tool takes a peptide sequence with at least 5 amino acid residues as input and computes the number of amino acids in it, its molecular weight, theoretical pI, extinction coefficient, amino-acid composition, half-life, aliphatic index, instability index, theoretical pI, grand average of hydropathicity (GRAVY) score, no. of positively charged residues, no. of negatively charged residues and atomic composition (Gasteiger et al. 2005).This tool applies N-end rule to estimate half-life, weight value of dipeptides to measure instability index, factors of relative volume occupied by aliphatic amino acids for aliphatic index and for GRAVY score, it calculates total of all hydropathy values of amino acids divided by the number of amino acids in input.

### Optimization of the construct and prediction of secondary structure

Different combinations of possible mutations in the construct were tried out using the Rational Design tool (http://www.camp.bicnirrh.res.in/predict_c21/) to reach desired physicochemical property scores calculated with ProtParam tool. Thus, an optimized sequence was designed. The secondary structure of this peptide was predicted with PSIPRED tool (http://bioinf.cs.ucl.ac.uk/psipred/). PSIPRED allots two feed-forward neural networks to execute the prediction. (Buchan and Jones 2019; Jones 1999).

### Prediction of antigenicity, allergenicity and toxicity of the final construct

The antigenicity of the final peptide was predicted with VaxiJen 2.0 tool (http://www.ddg-pharmfac.net/vaxijen/VaxiJen/VaxiJen.html). VaxiJen implements auto cross covariance (ACC) mediated transformation of amino acid sequence into the uniform vectors of prime properties of the amino acids (Doytchinova and Flower 2007, 2008). The allergenicity was determined using Allergen FP (http://ddg-pharmfac.net/AllergenFP/)and AllerTOP (https://www.ddg-pharmfac.net/AllerTOP/). AllergenFP also applies a similar method of generating vectors with ACC transformation (Dimitrov et al. 2014). The vectors are subsequently converted into binary fingerprints along with comparison in terms of Tanimoto coefficient. AllerTOP transforms protein sequences into uniform equal-length vectors applying auto cross covariance (ACC). Five E-descriptors ranging from amino acid hydrophobicity, β-strand forming propensity, helix-forming propensity to molecular size and relative abundance form the principal properties of the amino acids. The proteins are gradually categorized with k-nearest neighbor algorithm (kNN, k=1) relying on preknown set of 2427 allergens and 2427 non-allergens. The ToxinPred server (https://webs.iiitd.edu.in/raghava/toxinpred/index.html) was used to predict the toxicity of peptide candidate. It utilizes motif-based methods, dipeptide based Support Vector Machine (SVM) methods and hybrid prediction models based on toxic as well as non-toxic peptide datasets to generate predictions (Gupta et al. 2013).

### Identification of bacterial targets of the designed peptide

The tool of the Database of Antimicrobial Activity and Structure of Peptide (DBAASP) v3.0 (https://dbaasp.org/prediction/special) was used to predict peptide activity against specific bacterium. The tool predicts activity of the query peptide against microorganisms on the basis of existing information on bioactivity, toxicity and tertiary structure of more than 15,700 AMPs in DBAASP (Pirtskhalava et al. 2021). This tool confirms a peptide to be active with positive predictive value and MIC value less than 25 μg/ml while negative predictive value and MIC greater than 100 μg/ml for non-active ones. Then, the antibacTR tool (http://bioinf.uab.cat/antibactr) and the DrugBank (https://go.drugbank.com/) were used to find the druggable targets in the identified bacteria from DBAASP. The antibacTR expedites the essential prerequisite step of antibacterial drug discovery i.e., identification of potential antibacterial targets from the database of fully sequenced Gram-negative pathogens (Panjkovich et al. 2014). The ranking provided for each of the protein smoothens the task to discern a potential target. The DrugBank includes comprehensive details on drugs and drug targets.

### Prediction of possible human receptor targets of the peptide

After prediction of prokaryotic targets, it was essential to predict the human receptors of the designed peptide to estimate its therapeutic applications in the human body. So, probable receptors were identified with Swiss Target Prediction tool (http://swisstargetprediction.ch/). This tool executes a prediction algorithm involving principles of similarities of input molecule in 2D and 3D aspects with a pre-existing library of large no. of known active sites on several proteins in human (Daina et al. 2019).

### Selection of library of existing drugs against identified human receptors

The drugs widely used against the identified receptors were to be listed since they would act as controls for the designed test peptide. Scientific literatures and online databases like ChEMBL (https://www.ebi.ac.uk/chembl/), PubChem (https://pubchem.ncbi.nlm.nih.gov/) were the noteworthy resources used to complete the list (Gaulton et al. 2012; Kim et al. 2021).

### Tertiary structure modelling and structure assessment of the construct

It is essential to predict the stable conformation of the peptide drug. So, the I-TASSER server (https://zhanggroup.org/I-TASSER/) was used to model the tertiary structure of the peptide. The Structure Assessment Swiss (https://swissmodel.expasy.org/assess), ERRAT (https://servicesn.mbi.ucla.edu/ERRAT/) and ProSA-web servers (https://prosa.services.came.sbg.ac.at/prosa.php) were used to evaluate the structures. Ramachandran plot was also generated for the structure. The I-TASSER generates models based on PDB based templates identified with a multiple threading approach in Local Meta-Threading Server (LOMETS), and iterative assembly of fragments generated from threading templates. Moreover, this tool estimates the global accuracy as well as residue specific global quality of models (Yang and Zhang 2015). ERRAT analyses the erroneous segments in the structures with the pattern of nonbonded interactions (Sippl 1993). ProSA-web highlights the quality scores and subsequent flaws in the structures (Wiederstein and Sippl 2007). The Structure Assessment provides structural information and generates Ramachandran plot (Kiefer et al. 2009).

### Refinement and validation of the crude model

The global and local structure qualities of server generated model were improved with GalaxyRefine (http://galaxy.seoklab.org/refine) tool. The refined structure was validated again with structure assessment servers ERRAT and ProSA-web. The GalaxyRefine rebuilds sidechains followed by repackaging of sidechain leading to overall structure relaxation (Heo et al. 2013).

### Molecular docking studies

The protein model was computationally docked to all the identified targets in *Escherichia coli* and human receptors using the Patchdock (https://bioinfo3d.cs.tau.ac.il/PatchDock/) online server. The solved 3D structures of the target receptors were accessed from PDB. Patchdock applies a faster molecular docking algorithm based on matching of complementary patches. It applies geometric hashing and pose clustering to accelerate the computational processing time. Moreover, its rapid transformational search due to local feature matching leads to high docking efficiency (Duhovny et al. 2002, 2005). The obtained docking interactions were refined with Firedock (http://bioinfo3d.cs.tau.ac.il/FireDock/). The Firedock performs a high throughput refinement on protein-protein docked complexes applying side-chain optimization. It gradually provides scoring of docked solutions obtained from fast rigid-body docking (Andrusier et al. 2007; Mashiach et al. 2008). The Patchdock as well as Firedock scores were compared for each of the docking experiments performed. Similarly, all the existing drug molecules were docked with respective receptors as control docking experiments for human receptors.

### Interaction studies

In the cases of docking experiments with targets of *Escherichia coli*, *Klebsiella pneumoniae* and human, the complexes which had Patchdock (PD) and Firedock scores (FD) considerably higher than others were further selected for interaction study with Ligplot^+^ and PDBePISA (https://www.ebi.ac.uk/msd-srv/prot_int/cgi-bin/piserver). The interaction sites and residues were visualized with Pymol visualization software. Ligplot+ generates 2D interaction maps between protein and ligand (Laskowski and Swindells 2011). PDBePISA executes the analysis of the binding affinity parameters, interface study, residue interactions in binding partners of macromolecular complexes (Krissinel and Henrick 2007).

### Study of conformational dynamics of chosen docked complexes

The complexes of peptide with human and bacterial targets having best docking scores were subjected to conformational dynamics studies with the iMODS server (http://imods.chaconlab.org/). This tool implements Normal Mode Analysis (NMA) for computing internal coordinates and reproduces the probable trajectories of conformational changes of input macromolecules. It provides information on deformability of protein structure, eigen values and B-factor depicting disorder in protein atoms, to estimate the stability of macromolecules. It operates with rapid solutions to eigen problems extending the analysis space to larger macromolecules and applies an affine-model approach to simplify visualization of normal modes (López-Blanco et al. 2014).

## Results

### Preparation of the peptide construct

The AMPs found from the database belonged to Archaebacteria *Methanosarcina acetovorans* C2A and *Sulfolobus acidocaldarius* DSM 639. Selected sequences ‘LLELLGTPNNPGNVFKSNTL’ in *Methanosarcina acetovorans* C2A and ‘IIVSPLIKGLAVVSKKGFYV’ in *Sulfolobus acidocaldarius* DSM 639 have common Linocin M18 antibacterial motif (Fig. 1). These two peptides were joined with GGSG linker to form ‘LLELLGTPNNPGNVFKSNTGGSGIIVSPLIKGLAVVSKKGFYV’.

### Physicochemical properties and secondary structure of the construct

The physicochemical properties were computed for initial construct and using the Rational design tool along with mutations, new construct with optimum property scores was prepared. The initial designed sequence was provided as input in the Rational Design tool with default parameters of AMP probability result sorting option, 100 best hits and SVM algorithm. The obtained construct was found to have optimum physicochemical properties. Comparison of obtained scores for the physicochemical properties of respective constructs is shown in Table 1. The sequence of final construct is ‘LKELLGTPNNPGNVKKSNTLGGSGIIVSPLIKGLAKVSKKGFYK’ (Fig. 2). PSIPRED predicted secondary structure of peptide had 35% alpha-helix, 5% sheet and 60% coiled structure (Fig. 3).

**Table 1.**
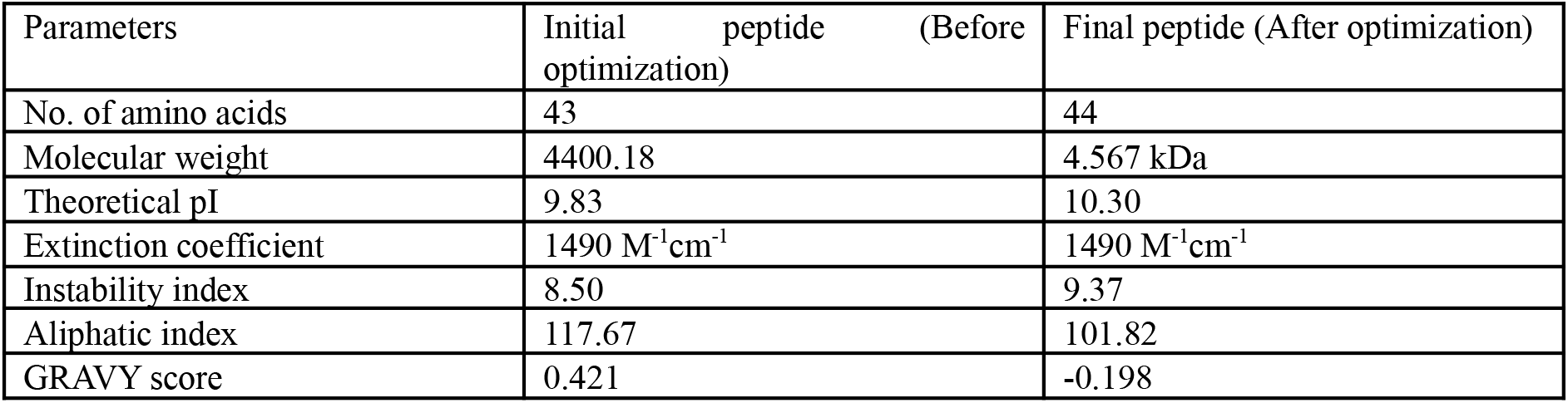
Peptide properties.

| Parameters | Initial peptide (Before optimization) | Final peptide (After optimization) |
| --- | --- | --- |
| No. of amino acids | 43 | 44 |
| Molecular weight | 4400.18 | 4.567 kDa |
| Theoretical pI | 9.83 | 10.30 |
| Extinction coefficient | 1490 M<sup>-1</sup>cm<sup>-1</sup> | 1490 M<sup>-1</sup>cm<sup>-1</sup> |
| Instability index | 8.50 | 9.37 |
| Aliphatic index | 117.67 | 101.82 |
| GRAVY score | 0.421 | -0.198 |

**Fig. 2.**
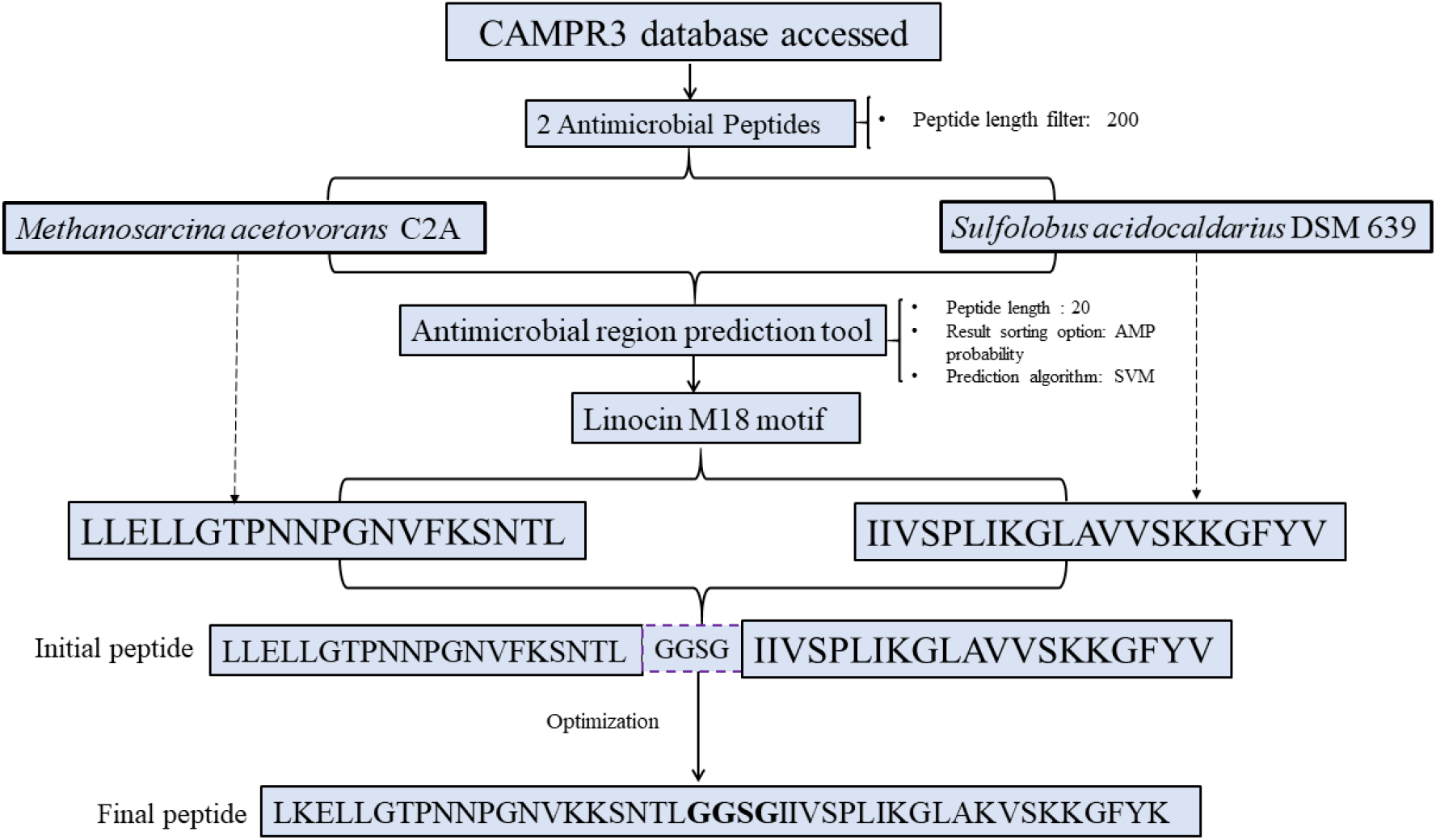
Schematic diagram of the designing strategy of peptide construct

**Fig. 3.**
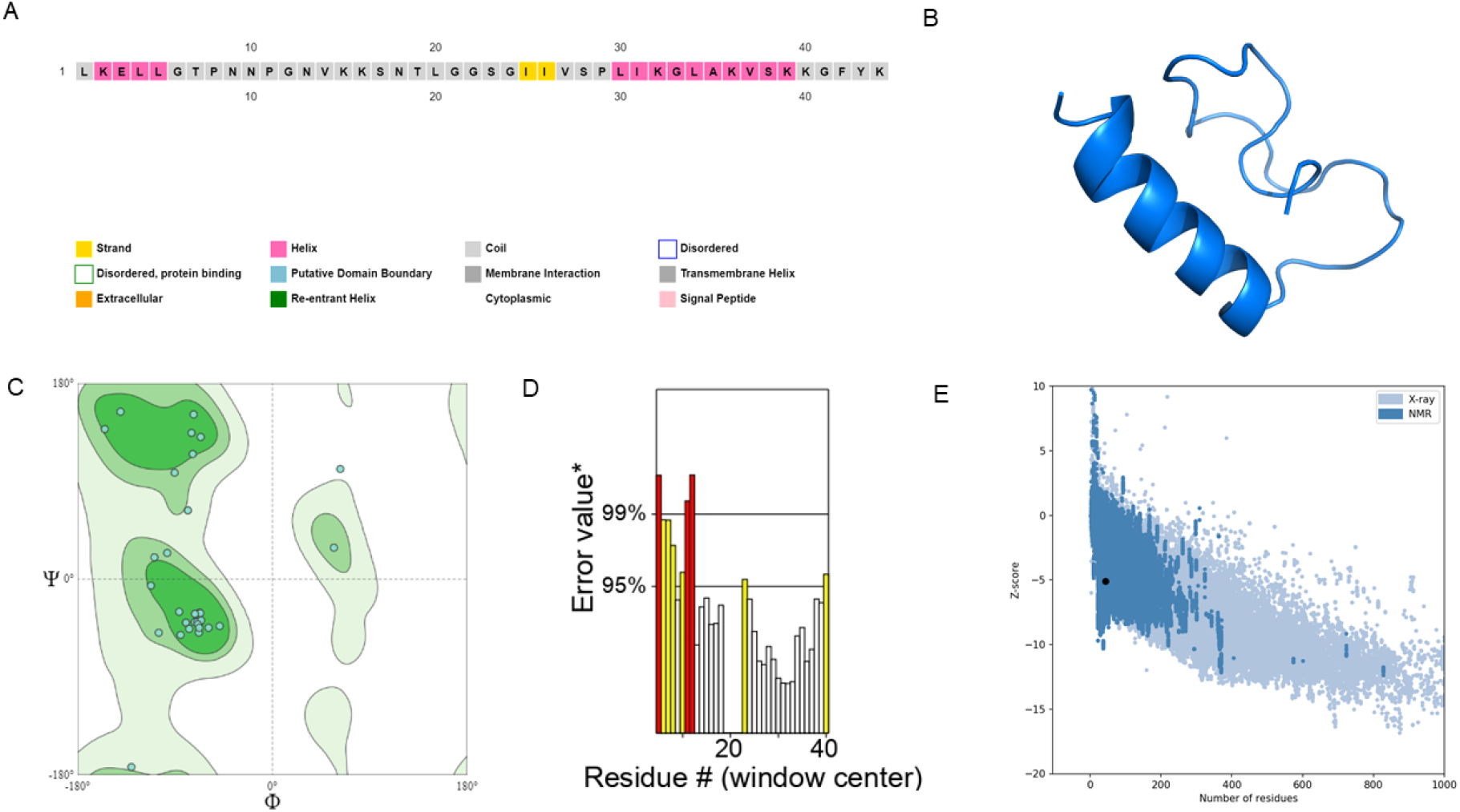
Peptide secondary structure prediction and modelling its tertiary structure along with model evaluation **a** PSIPRED predicted secondary structure. **b** Refined structure of I-TASSER generated tertiary model. **c** Ramachandran plot generated for the model from Swiss Structure Assessment server. **d** Model evaluation plot generated with ERRAT **e** Z score assessment of the model (marked as black dot) from ProSA-web

### Antigenicity, allergenicity and toxicity of the peptide

One of the primary criteria in peptide drug therapy is its non-toxic characteristic inside the human body. The VaxiJen 2.0 results obtained for tumour as target and 0.4 as threshold estimated peptide to be probable antigen. AllergenFP and AllerTOP servers confirmed the peptide to be non-allergic. ToxinPred predicted the peptide to be non-toxic with default parameters and SVM based prediction method as input.

### Bacterial targets

The peptide was found to strongly bind *Escherichia coli* and *Klebsiella pneumoniae* targets. Computational predictions estimated druggable targets in the bacteria. Only those targets with solved tertiary structures were considered. The targets selected in *Escherichia coli* were 30S ribosomal protein S18, Inorganic pyrophosphatase, Flavodoxin −1, 2,3,4,5 – tetrahydropyridine-2,6-dicarboxylate N-succinyltransferase, 30S ribosomal protein S4, DNA directed RNA polymerase subunit α, UDP-3-O-[3-hydroxymyristoyl] N-acetylglucosamine deacetylase and 3-hydroxydecanoyl-[acyl-carrier-protein] dehydratase and in *Klebsiella pneumoniae*, acetolactate synthase, metallo-beta lactamase, carbapenemase and SHV-1 beta-lactamase were identified.

### Human receptor targets

Cathepsin D (CTSD), Caspase-1 (CASP1), Signal transducer and activator of transcription-3 (STAT3), Beta-secretase 2 (BACE1), Beta secretase 1 (BACE2), Renin, SRC, Mitogen activated protein kinase-1 (MAPK1), Angiotensin-converting enzyme (ACE), Angiotensin-converting enzyme 2 (ACE2), 3-hydroxy-3-methyl glutaryl coenzyme A (HMG CoA) and Ribonucleotide reductase (RNR) were identified as the possible targets in SwissTargetPrediction tool for the peptide candidate. Majority of these targets play significant role in various diseases like renal fibrosis, dysregulation of apoptosis, cardiovascular defects, hypertension, neuronal degeneration, different carcinomas like pancreatic cancer, lymphoma, glioma etc (Fox et al. 2016, Knight and Barrett 1976; MacKenzie et al. 2010; Li et al. 2019; Chang et al. 2017, Abletshauser et al. 2002; Jackson et al. 1984; Keikhaei et al. 2016; Aye et al. 2015; Farris et al 2021).

### Existing drug candidates against the human receptors

Evidences from available literatures confirmed existing drugs against the receptors identified. Pepstatin inhibits Cathepsin-D (Knight and Barrett 1976; Fox et al. 2016), belnacasan and z-yad-fmk are inhibitors of Caspase-1 (MacKenzie et al. 2010; Chopra et al. 2009), crytotanshinone, niclosamide and napabucasin inhibit STAT3 (Li et al. 2019, 2013, 2020), Phenserine inhibits BACE2 (Chang et al. 2017), BACE1 inhibited by elenbecestat and umibecestat (Adewole and Ishola 2021; Tran et al. 2020), aliskiren strongly binds and inhibits Renin (Wal et al. 2011), SRC inhibited by kx-2391, bosutinib and saracatinib (Naing et al. 2013, Vultur et al. 2008; Gucalp et al. 2011), ulixertinib and ravoxertinib inhibit MAPK1 (Braicu et al. 2019), enalapril, benazepril inhibit ACE (Jackson et al. 1984; Yu et al. 2006), hydroxychloroquine inhibits ACE2 (Fu et al. 2021), HMG-CoA, rosuvastatin and simavastatin targeted against HMG-CoA (Abletshauser et al. 2002; Slater and MacDonald 1988) and hydroxyurea inhibits RDR (Keikhaei et al. 2016; Singh and Xu 2016).

### Modelled tertiary structure of the construct and validation of refined model

The tertiary structure of the peptide was modelled using I-TASSER server. I-TASSER provided top 5 models and the model with highest C score (−2.37) was selected since it depicts good quality of the model. Then, GalaxyRefine was used to refine the model and improve the quality assessed by Ramachandran plots, ERRAT and ProSA-web. Model-2 with optimum RMSD (0.523), GDT-HA (0.9432), MolProbity (2.992), Clash score (21.7), poor rotamers (5.6) among the five models generated by GalaxyRefine server, was selected. It showed promising results in Ramachandran plot where 96.875% residues were found to be in the favoured region while 3.125% residues in the allowed region. The ERRAT score of 71.875 (good models have values greater than 50) and −5.1 (z score) from ProSA-web bolstered the reliability and improved quality of the predicted protein model (Fig. 3).

### Preparation of receptors for molecular docking

The solved tertiary structures of *Escherichia coli*, *Klebsiella pneumoniae* and human receptors were accessed from PDB for the purpose of molecular docking with the peptide. Any of the bound ligands or inhibitors in receptor structures were removed before docking. The PDB ID of respective *Escherichia coli* targets are: Flavodoxin 1: 1AG9, UDP-3-O-[3-hydroxymyristoyl] N-acetylglucosamine deacetylase: 6P89, 3-hydroxydecanoyl-[acyl-carrier-protein] dehydratase: 4KEH, DNA-directed RNA polymerase subunit α: 3K4G, 30S ribosomal S4: 6XZB, Inorganic pyrophosphatase: 1FAJ, 30S ribosomal S18: 3JCE. PDB ID of *Klebsiella pneumoniae* targets are acetolactate synthase: 1OZF, carabapenemase: 2OV5, SHV-1 betalactamase: 1ONG, metallo-beta lactamase: 3PG4. PDB ID of respective human receptors: CTSD: 4OBZ, CASP1: 1BMQ, STAT3: 4ZIA, BACE2: 2EWY, BACE1: 1FKN, RENIN: 2V07, SRC: 1A07, MAPK1: 4ZXT, ACE: 1O8A, ACE2: 1R42, HMG: 1DQ8, RNR: 2WGH.

### Molecular docking

The computational docking of refined peptide model and various targets, performed in PatchDock tool followed by docking refinement with FireDock tool provided docking scores and global energy scores respectively. The targets showing high PatchDock scores and low FireDock global energy scores were selected for further visualization of complex and interacting residues. The CTSD, BACE-2 and RNR were chosen amongst the other human receptors. The docking scores of these targets were even better than their existing drug counterparts as shown in Table 2. Flavodoxin-1, UDP-3-O-[3-hydroxymyristoyl] N-acetylglucosamine deacetylase and 3-hydroxydecanoyl-[acyl-carrier-protein] dehydratase from *Escherichia coli* and acetolactate synthase of *Klebsiella pneumoniae* were selected on the basis of scores as displayed in Table 3. The detailed docking scores of all docked complexes are mentioned in Supplementary Table 1, 2 and 3.

**Table 2.** Docking scores for peptide-human receptor complexes.

| SL NO | Receptor | PEPTIDE docking score |  | Existing candidate drug | DRUG Docking score |  |
| --- | --- | --- | --- | --- | --- | --- |
|  |  | PATCH DOCK | FIRE DOCK energy score |  | PATCH DOCK | FIRE DOCK energy score |
| 1. | CTSD | 13974 | -55.81 | Pepstatin | 7334 | -49.04 |
| 2. | BACE2 | 14030 | -59.55 | Phenserine | 5444 | -48.57 |
|  |  |  |  | Posiphen | 5460 | -53.31 |
| 3. | RNR | 15406 | -35.07 | Hydroxyurea | 1502 | -12.92 |

**Table 3.** Docking scores for peptide-bacterial target complexes.

| Organism | Sl NO. | Receptor | PEPTIDE docking score |  |
| --- | --- | --- | --- | --- |
|  |  |  | PATCH DOCK | FIRE DOCK energy score |
| *Escherichia coli* | 1. | Flavodoxin 1 | 12704 | -40.36 |
|  | 2. | UDP-3-O-[3-hydroxymyristoyl] N-acetylglucosamine deacetylase | 13580 | -51.95 |
|  | 3. | 3-hydroxydecanoyl-[acyl-carrier-protein] dehydratase | 12272 | -39.66 |
| *Klebsiella pneumoniae* | 1. | Acetolactate synthase | 12956 | -58.61 |

### Interactions in selected docked complexes

The selected receptors were further subjected to interaction mapping at the peptide-target binding interface with computational methods. The residues involved in strong hydrogen bond interactions in docked complexes with bacterial and human receptors have been listed in Table 4. The peptide interaction site in the docked complexes of bacterial targets (Fig.4) and human receptors (Fig. 5) with highest docking scores have been visualized with molecular visualization tool.

**Table 4.** Residues involved in hydrogen bonding interactions in selected docked complexes.

| ORGANISM | RECEPTOR | PEPTIDE |
| --- | --- | --- |
| *Escherichia coli* | Flavodoxin-1:<br>Asp 77, Arg 112, Thr 72, Glu 61,<br>Glu 75 | Asn 18, Ser 28, Lys 15, Val 14 |
|  | UDP-3-O-[3-hydroxymyristoyl]<br>N-acetylglucosamine deacetylase:<br>Glu 49 | Asn 9 |
|  | 3-hydroxydecanoyl-[acyl-carrier-protein] dehydratase:<br>Asn 48, Lys 112, Gln 14, Glu 49,<br>Glu 20, Glu 110, Ser 158 | Asn 9, Thr 19, Asn 18, Ser 28,<br>Tyr 43, Gly 21 |
| *Klebsiella pneumoniae* | Acetolactate synthase:<br>Glu 310, Ser 121 | Lys 39, Gly 21, Gly 22 |
| Human | CTSD:<br>His 57, Cys 27, Trp 54 | Thr 7, Gly 21, Lys 16 |
|  | BACE2:<br>Asp 25, Lys 130, Asn 183, Pro<br>321, Ala 173, Gln 23 | Lys 36, Tyr 43, Asn 18, Ser 28,<br>Val 14, |
|  | RNR:<br>Lys 316, Arg 284, Thr 741, Glu<br>321, Lys 320, Glu 322 | Lys 2, Gly 22, Asn 9, Asn 10,<br>Pro 11 |

**Fig. 4.**
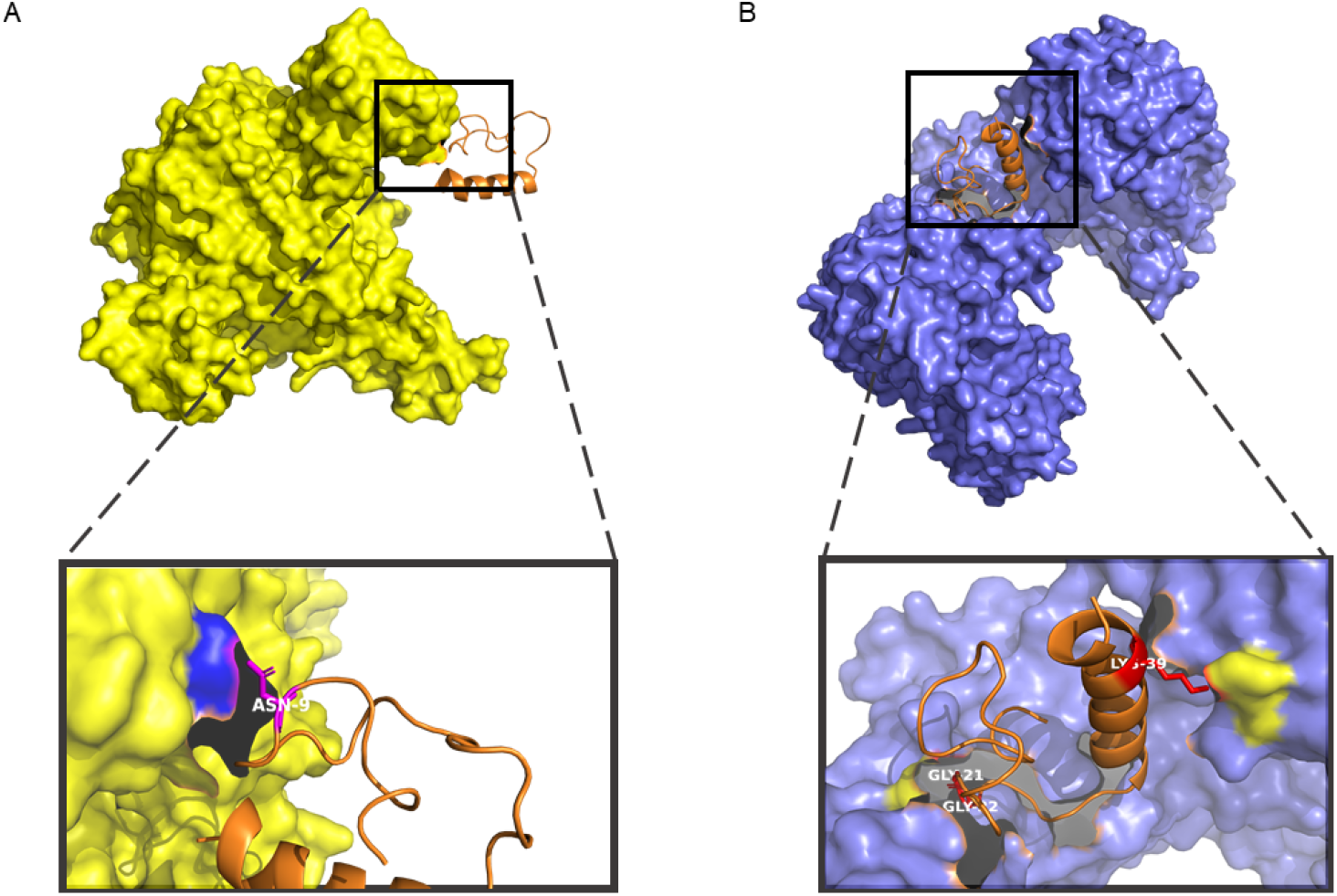
Visualization of interactions in the docked complexes of peptide (orange) with **a** UDP-3-O-[3-hydroxymyristoyl] N-acetylglucosamine deacetylase (yellow) of *Escherichia coli* and **b** acetolactate synthase (ultramarine blue) of *Klebsiella pneumoniae*. Interacting residues of peptide are labelled white and that of the receptor represented with colored surfaces.the receptor represented with yellow-colored surfaces.

**Fig. 5.**
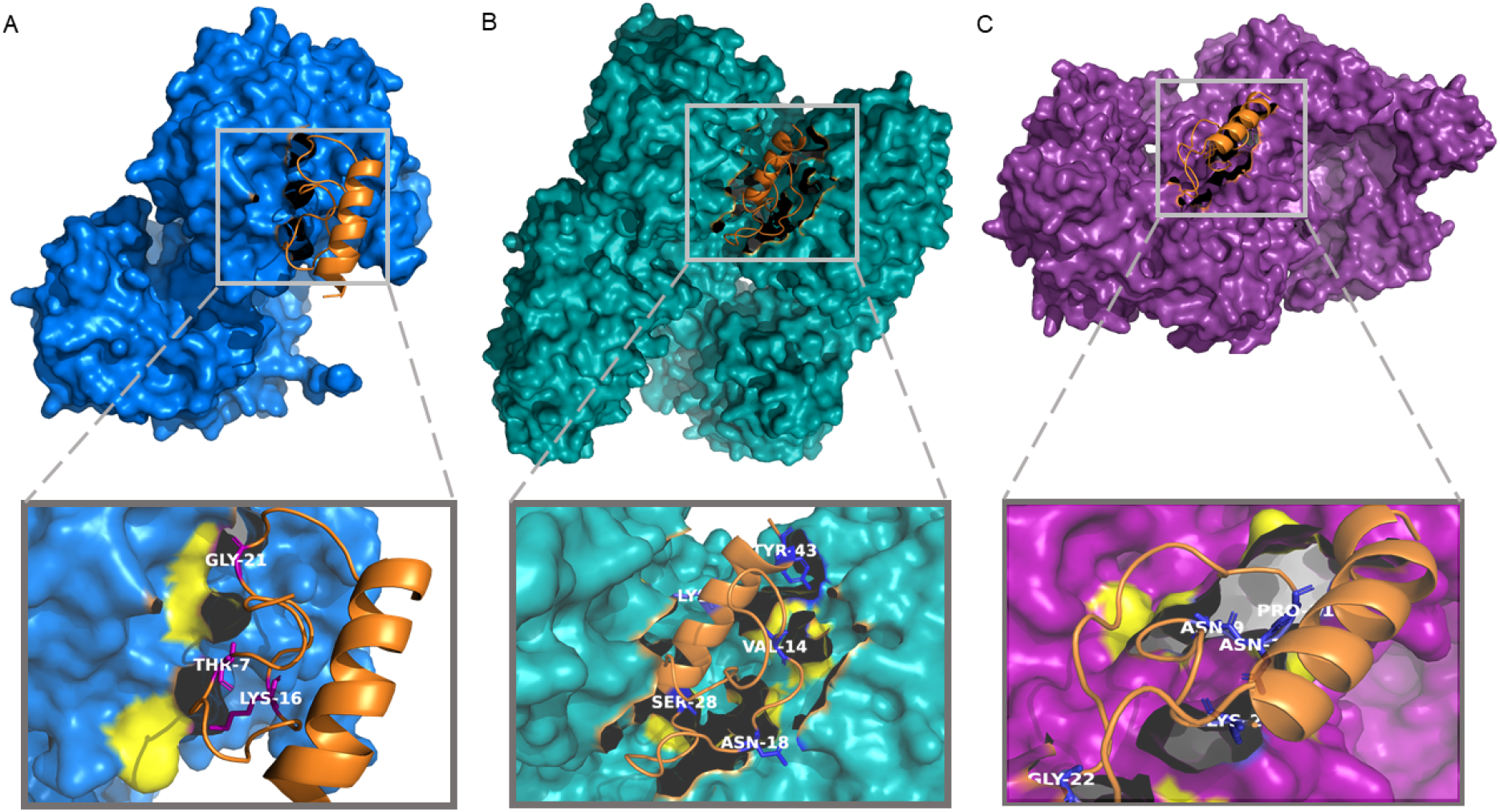
3D visualization of interactions in the docked complexes of peptide (orange) with **a** Cathepsin D (blue) **b** BACE2 (deep green) **c** RNR (purple). Interacting residues of peptide are labelled white and that of the receptor represented with yellow-colored surfaces.

### Prediction of complex dynamics from iMODS

The iMODS server predicted stability as well as flexibility of selected docked complexes. The deformability, NMA computed B-factor (NMB) and eigen values were computed for the complexes.

#### *Escherichia coli* target docked complexes

Flavodoxin-peptide complex showed overall high deformability in the main chain in comparison to other targets (Suppl. Fig. 2a). The UDP-3-O-[3-hydroxymyristoyl] N-acetylglucosamine deacetylase showed lowest deformability except some spikes in few internal and C-terminal stretches (Fig. 6a). It has least no. of hinges in main chain than others. The NMA calculated B-factors indicate the extent of distortions at the atomic level. The NMA B factor (NMB) was minimized in comparison to PDB B factor for UDP-3-O-[3-hydroxymyristoyl] N-acetylglucosamine deacetylase (Fig. 6b). while the NMA B-factor was greater than B-factor of PDB (BPB) to a great extent with some exceptional atomic stretches in Flavodoxin-peptide complex (Suppl. Fig. 2d). The NMB for 3-hydroxydecanoyl-[acyl-carrier-protein] dehydratase – peptide almost overlapped with BPB except a few regions (Suppl. Fig. 6e). The computed eigen values of the peptide docked with Flavodoxin-1, 3-hydroxydecanoyl-[acyl-carrier-protein] dehydratase and UDP-3-O-[3-hydroxymyristoyl] N-acetylglucosamine deacetylase were 1.288 x 10^−4^, 8.95 x 10^−5^ and 7.13 x 10^−6^ respectively (Suppl. Fig. 2). These values provide an estimation of the energy for structure deformation. So, it could be stated that Flavodoxin-peptide complex could not be deformed easily than others since it has the highest eigen value correlating to more energy requirement to deform its structure. Although 3-hydroxydecanoyl-[acyl-carrier-protein] dehydratase has a little higher eigen value (Suppl. Fig. 2h) than UDP-3-O-[3-hydroxymyristoyl] N-acetylglucosamine deacetylase, evaluating all the aspects of plots generated from iMODS, the latter could be stated to be involved in stable interactions with the peptide than other targets. So, it could be accepted as a suitable target on the whole.

**Fig. 6.**
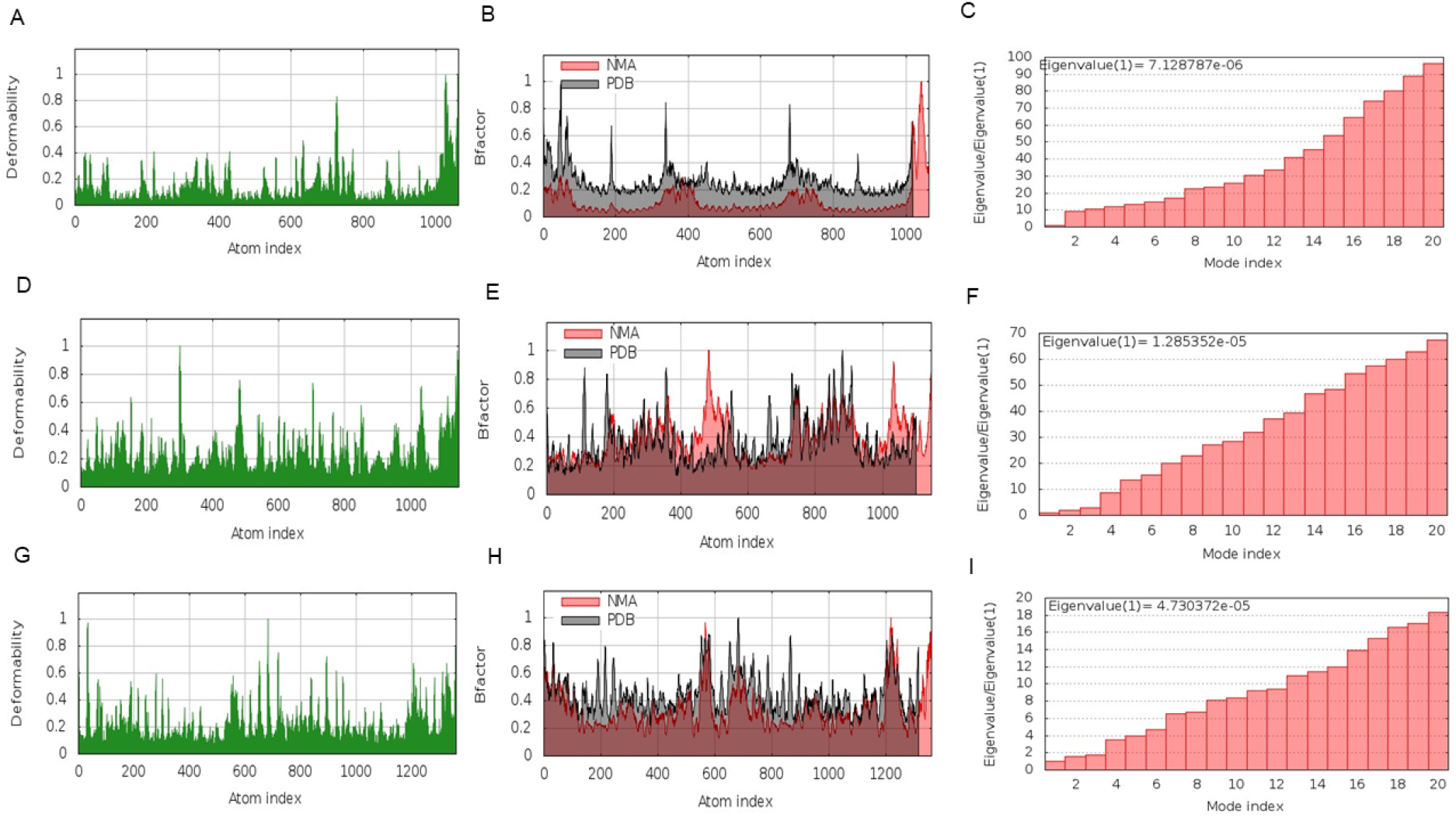
Predictions of conformational dynamics from iMODS. *Escherichia coli* UDP-3-O-[3-hydroxymyristoyl] N-acetylglucosamine deacetylase - peptide complex **a** deformability plot **b** B-factor calculations **c** Eigen value estimation. *Klebsiella pneumoniae* acetolactate synthase- peptide complex **d** deformability plot **e** B-factor calculations **f** Eigen value estimation. RNR-peptide complex **g** deformability plot **h** B-factor calculations **i** Eigen value estimation.

#### *Klebsiella pneumoniae* target docked complexes

The analysis of iMODS results for the peptide and acetolactate-synthase complex showed quite a low deformability in its main chain, NMA B-factor values overlapping to a greater extent with PDB B-factor except few internal and C-terminus regions (Fig. 6e). The eigen value obtained, 1.285 x 10^−5^, was even found to be pretty high confirming its stability (Fig. 6f). So, undeniably, this enzyme of Klebsiella pneumoniae could be a probable target for this peptide construct.

#### Human target docked complexes

The RNR-peptide complex showed least deformability among others indicating presence of less hinges in the main chain (Fig. 6g). It also showed minimized NMB values than PDB B-factor values (Fig. 6h). The BACE-2 complex showed highest deformability (Suppl Fig. 3) indicating presence of more hinges in the main chain. The eigen values of the peptide complexes with Cathepsin D, BACE 2 and RNR were 1.32 x 10^−5^, 3.19 x 10^−5^ and 4.73 x 10^−5^ respectively (Suppl Fig. 3). These values provide an estimation that the RNR-peptide complex requires highest energy among others to suffer deformation (Fig. 6f). So, having an overview of all the plots (Suppl Fig. 3), the RNR-peptide complex tends to be the most stable one. Certainly, RNR tends to be a suitable intracellular target.

## Discussions

Recent times have witnessed the augmented market use of peptides as apposite therapeutics over small molecule drugs, due to multitudinous reasons, one of them being their high selectivity due to presence of multiple points of contacts with targets. Albeit delivery aspect of peptides has squelched their usage for a long time, advances are being made by use of absorption advancers, enzyme inhibitors, PNPs, nano-emulsions, muco-adhesives, nano-micelles etc. in recent studies (Bruno et al 2013). The fact that peptides possess the flexibility of being transmuted into any form to target a broad spectrum of target molecules, gives them the endless scope to be applied to fields such as immunology, oncology, infectious diseases, MDR bacterial infections etc. Moreover, globally emerging multidrug-resistant (MDR) bacterial infections have challenged the existing models of treatment using traditional antibiotics (Li and Webster 2018, Torgerson and Mapp 2017). The current study eminently focusses on utilization of the plausible antimicrobial properties of Archaeal bacteriocins to construct a probable peptide model which has shown promising results against both human diseases and MDR bacterial infections.

Our final peptide construct (LKELLGTPNNPGNVKKSNTLGGSGIIVSPLIKGLAKVSKKGFYK) has significant antimicrobial properties since it has been derived from two antimicrobial motifs (LLELLGTPNNPGNVFKSNTL, IIVSPLIKGLAVVSKKGFYV) as obtained from CAMPR3 database. The construct showed optimum scores for physicochemical properties as predicted by ProtParam tool. In addition, the predicted antigenic, non-allergenic and non-toxic nature of this construct galvanizes the fact for its further experimental assessments as a peptide drug. The refined 3D model of improved quality obtained for the peptide showed promising docking results with targets Flavodoxin 1, UDP-3-O-[3-hydroxymyristoyl] N-acetylglucosamine deacetylase, 3-hydroxydecanoyl-[acyl-carrier-protein] dehydratase of *Escherichia coli*, acetolactase synthase of *Klebsiella pneumoniae* and human targets: CTSD, BACE-2 and RNR. The peptide was involved in strong hydrogen binding interactions showing high affinity to the mentioned receptors. Analyses from the iMODS server additionally verified high stability with low variance in the interactions of peptide with UDP-3-O-[3- hydroxymyristoyl] N-acetylglucosamine deacetylase, acetolactase synthase and RNR targets. The high affinity to these targets confers certain advantages. The deacetylase is a key player in the lipid A biosynthesis pathway in *Escherichia coli* (Sorensen et al 1996). So, blocking this enzyme with the peptide could be an appropriate measure to inhibit bacterial growth. Secondly, acetolactate synthase plays a significant role in butanediol fermentation by the bacterium (Pang et al 2002). In addition, Klebsiella pneumoniae is one of the noteworthy MDR bacterium lacking widely recognized treatment options (Bassetti et al 2018). So, blocking such enzyme could actually provide a possible measure to disrupt bacterial growth. Lastly, it is a well-known fact that cancer cells require RNR for synthesizing deoxyribonucleotide triphosphates (dNTP) de novo (Aye et al. 2015). RNR catalyzes the rate limiting step for all dNTP biosynthesis. So, targeting such an enzyme with this peptide could be an effective treatment against cancer. It has also been found that our peptide binds strongly to the RNR in comparison to its well-known inhibitor, hydroxyurea. Although the therapeutic spectrum of hydroxyurea reaches out to neoplastic (carcinomas) as well as non-neoplastic diseases (sickle cell anemia), the cytotoxic effects exerted by it even in normal cells due to prolonged usage gains limitations in its application (Singh and Xu 2016). Continuous usage of it has shown increasing risks of occurrence of skin cancers and other side-effects (Kerdoud et al 2021). So, our peptide could possibly be an efficient alternative to such traditional inhibitors. The peptide mediated inhibition of BACE-2 and CTSD could also be a beneficial therapy since BACE-2 plays a significant role in enhanced neurodegeneration in Alzheimer’s disease, tumor progression such as in glioma, melanoma etc. (Farris et al 2021) and CTSD is also involved in various neurodegenerative disorders and cancers such as melanoma, breast cancer, prostate cancer etc (Mijanovic et al 2021).

Considering about the targeted delivery for destruction of bacterial biofilms which can be omnipresent on medical implants, indwelling devices, wounds (Lebeaux et al 2013), future studies must be conducted in wet laboratory, developing our peptide model into a viable peptide drug and combining it with existing PNPs, biodegradable polymer-stabilized oil-in-water nano-sponges (BNS) (Nabawy et al. 2020) to construct bioconjugates and checking their minimal inhibitory concentrations (MIC), fractional inhibitory concentration index (FICI), synergistic interactions, additive interactions (Chen and Zhong 2017, Gupta et al. 2020) to confirm their efficacy against extracellular polymeric substances and bacterial colonies. Recent studies suggest the development of bioconjugates between polymeric nanoparticles (PNPs) and existing antibiotics has been proved to degrade the biofilms and augment the efficacy of the drug against the bacteria (Tew et al 2010; Gupta et al 2019). These PNPs not only assure the targeted delivery of the drug in the bacterial colony, but simultaneously also enhances the antibiotic potency at a very low dosage. These techniques of combination therapy ought to be validated experimentally with our peptide model. Even the strong affinity of this stable, non-toxic, non-allergenic peptide construct towards human targets could undoubtedly be a standpoint for the initiation of peptide drug discovery. Moreover, future experimental studies with insights to systemic stability of peptide and site-specific delivery strategies via oral or transdermal routes are inevitable for validating its therapeutic potency against human targets (Bruno et al 2013).

## Conclusion

Bacterial biofilm development with time has made microbes to become unassailable by multiple existing antibiotics, which in turn has led to sundry chronic infections to become ubiquitous among us. This thorough yet compendious in silico study regarding the merits of archaeosins has shown promising results regarding the methodology and efficacies of peptide drug therapeutics involving AMPs. Using relative receptor-ligand docking studies, and computational in vivo stability characterization, it has been successfully shown that our designed peptide candidate can aptly act against well known targets associated with chronic human diseases and even against bacteria. All these in-silico predictions can be further corroborated using in vitro and in vivo experimental evidence ultimately leading to utilization of this archaeosin-peptide-therapeutic construct in pragmatic designing of a new broad-spectrum drug in future.

## Declarations

### Availability of data and material

The datasets generated during and/or analysed during the current study are available from the corresponding author on reasonable request.

### Author’s contributions

The authors have contributed equally to this work.

## Supplementary information

Supplementary Table 1 Docking scores for peptide-human receptor docked complexes. Supplementary Table 2: Molecular docking scores of all docked complexes of Escherichia coli targets with peptide. Supplementary Table 3: Molecular docking scores of all docked complexes of Klebsiella pneumoniae targets with peptide. Supplementary Fig. 1 Visualization of interactions in the docked complexes. Supplementary Fig. 2 Predictions of conformational dynamics from iMODS. Supplementary Fig. 3 Predicted conformational dynamics from iMODS.

