## Supplementary Information for "In silico design of bioactive chimeric peptide from archaeal antimicrobial peptides"

### Designing peptide drug with archaeal antimicrobial peptides against multidrug resistant bacterial biofilms and human diseases: an in silico approach

International Journal of Peptide Research and Therapeutics

Souvik Banerjee[1*], Soham Chakraborty[2], Kaustav Majumder[3]

[1]Department of Microbiology, St. Xavier's College (Autonomous), Kolkata, West Bengal, India

[2]Department of Chemistry, Indian Institute of Technology, Indian School of Mines (ISM), Dhanbad, Jharkhand, India

[3]Department of Biosciences and Bioengineering, Indian Institute of Technology, Bombay, Maharashtra, India

*Corresponding author

**Supplementary Table 1 Docking scores for peptide-human receptor docked complexes**

| SL NO. | Receptor | PEPTIDE docking score | | Existing drug candidate | DRUG Docking score | |
|---|---|---|---|---|---|---|
| | | PATCH DOCK | FIRE DOCK energy score | | PATCH DOCK | FIRE DOCK energy score |
| 1. | CTSD | 13974 | -55.81 | PEPSTATIN | 7334 | -49.04 |
| 2. | CASP1 | 10912 | -28.46 | BELNACASAN | 5168 | -42.17 |
| | | | | z-yad-fmk | 5940 | -33.8 |
| 3. | STAT3 | 13088 | -25.47 | CRYTOTANSHINONE | 4232 | -38.72 |
| | | | | NICLOSAMIDE | 4092 | -35.27 |
| | | | | NAPABUCASIN | 3622 | -27.34 |
| 4. | BACE2 | 14030 | -59.55 | PHENSERINE | 5444 | -48.57 |
| | | | | POSIPHEN | 5460 | -53.31 |
| 5. | BACE1 | 13326 | -21.56 | ELENBECESTAT | 6100 | -47.84 |

| | | | | | | |
|---|---|---|---|---|---|---|
| | | | | UMIBECESTAT | 6228 | -41.82 |
| 6. | RENIN | 12042 | -42.80 | ALISKIREN | 7534 | -58.49 |
| 7. | SRC | 11228 | -22.33 | Kx2-391 | 5688 | -39.38 |
| | | | | BOSUTINIB | 5348 | -34.51 |
| | | | | SARACATINIB | 5692 | -39.50 |
| 8. | MAPK1 | 11892 | -17.15 | ULIXERTINIB | 4992 | -39.55 |
| | | | | RAVOXERTINIB | 5494 | -41.27 |
| 9. | ACE | 10776 | -16.51 | ENALAPRIL | 5360 | -34.65 |
| | | | | BENAZEPRIL | 6278 | -44.04 |
| 10. | ACE2 | 15262 | -1.11 | HYDROXYCHLOROQUINE | 4796 | -37.04 |
| 11. | HMG | 13326 | -21.56 | HMG-CoA | 7758 | -47.58 |
| | | | | SIMAVASTATIN | 6168 | -42.97 |
| | | | | ROSUVASTATIN | 6170 | -43.52 |
| 12. | RNR | 15406 | -35.07 | HYDROXYUREA | 1502 | -12.92 |

**Supplementary Table 2: Molecular docking scores of all docked complexes of *Escherichia coli* targets with peptide**

| SL NO. | Receptor | PEPTIDE docking score | |
|---|---|---|---|
| | | PATCH DOCK | FIRE DOCK energy score |
| 1. | 30S ribosomal S18 | 9064 | -15.58 |
| 2. | Inorganic pyrophosphatase | 11218 | -20 |
| 3. | Flavodoxin 1 | 12704 | -40.36 |
| 4. | 30S ribosomal S4 | 11568 | -17.13 |
| 5. | DNA-directed RNA polymerase subunit α | 13540 | -31.95 |
| 6. | UDP-3-O-[3-hydroxymyristoyl] N-acetylglucosamine deacetylase | 13580 | -51.95 |
| 7. | 3-hydroxydecanoyl-[acyl-carrier-protein] dehydratase | 12272 | -39.66 |

**Supplementary Table 3: Molecular docking scores of all docked complexes of *Klebsiella pneumoniae* targets with peptide**

| SL NO. | Receptor | PEPTIDE docking score | |
|---|---|---|---|
| | | PATCH DOCK | FIRE DOCK global energy score |
| 1. | Acetolactate synthase | 12956 | -58.61 |
| 2. | Metallo-beta lactamase | 10478 | -37.22 |
| 3. | Carbapenemase | 14684 | -8.95 |
| 4. | SHV-1 beta-lactamase | 10538 | -14.84 |

**Supplementary figures:**

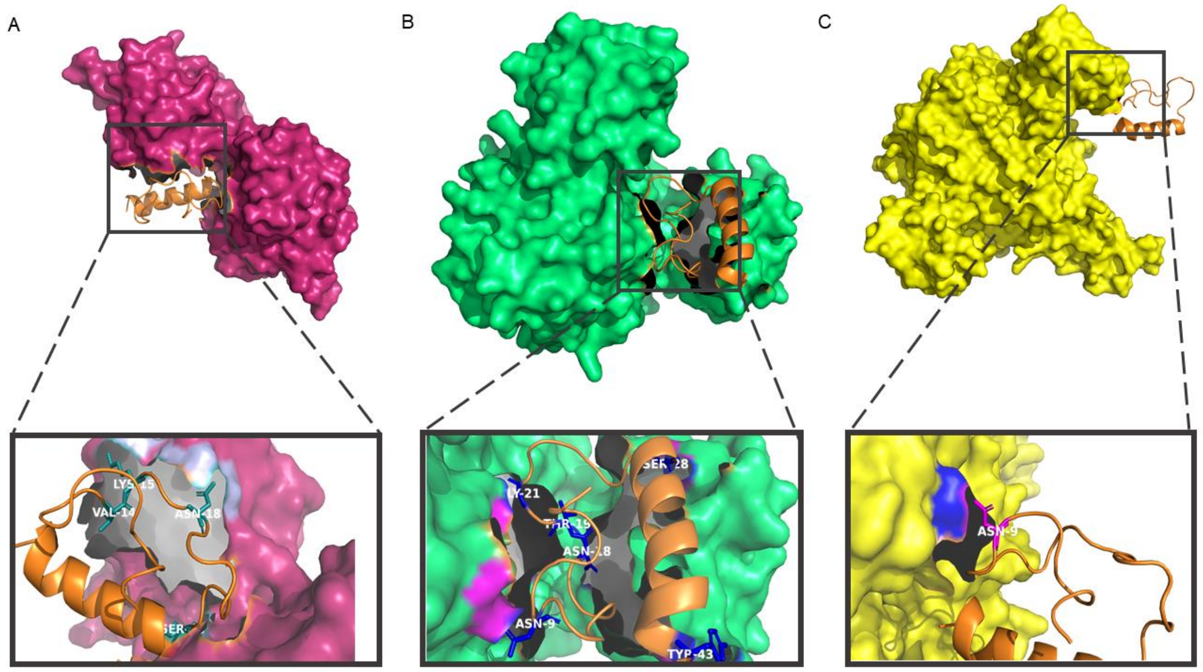


**Supplementary Fig. 1** Visualization of interactions in the docked complexes of peptide (orange) with **a** Flavodoxin 1 **b** 3-hydroxydecanoyl-[acyl-carrier-protein] dehydratase **c** UDP-3-O-[3-hydroxymyristoyl] N-acetylglucosamine deacetylase. Interacting residues labelled in white and receptor interface represented with surface.

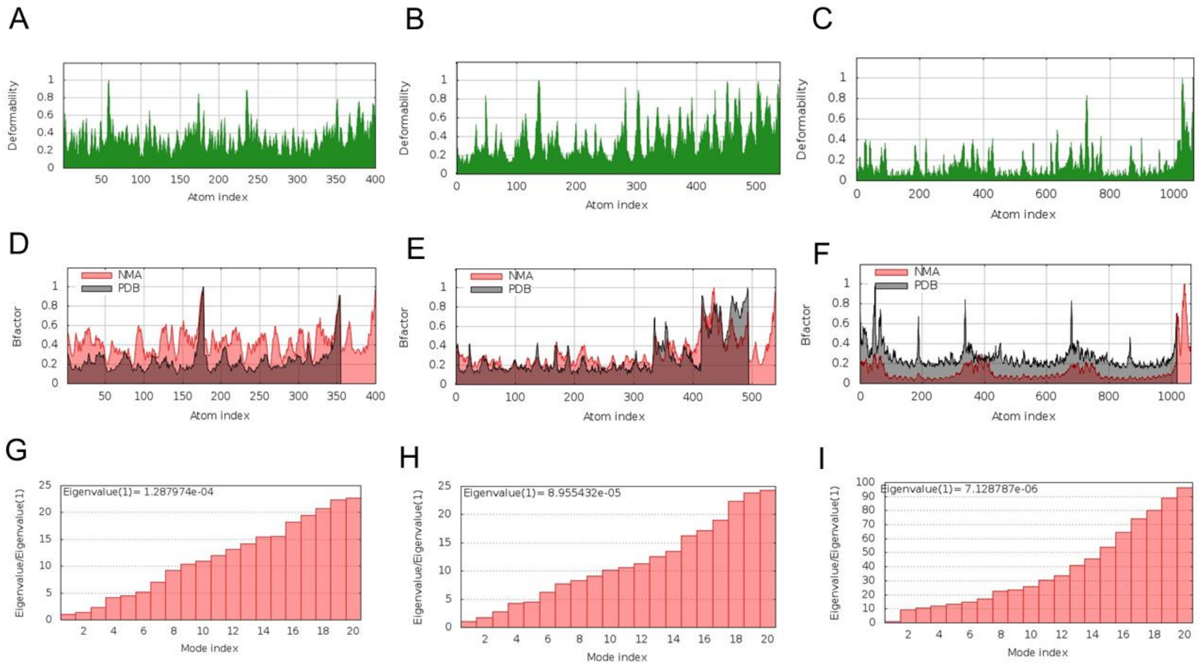

**Supplementary Fig. 2** Predictions of conformational dynamics from iMODS. Deformability plot for peptide complex with **a** Flavodoxin-1 **b** 3-hydroxydecanoyl-[acyl-carrier-protein] dehydratase **c** UDP-3-O-[3-hydroxymyristoyl] N-acetylglucosamine deacetylase. B-factor comparison for peptide complex with target **d** Flavodoxin-1 **e** 3-hydroxydecanoyl-[acyl-carrier-protein] dehydratase **f** UDP-3-O-[3-hydroxymyristoyl] N-acetylglucosamine deacetylase. Eigen-values for docked complex of peptide with **g** Flavodoxin-1 **h** 3-hydroxydecanoyl-[acyl-carrier-protein] dehydratase **i** UDP-3-O-[3-hydroxymyristoyl] N-acetylglucosamine deacetylase

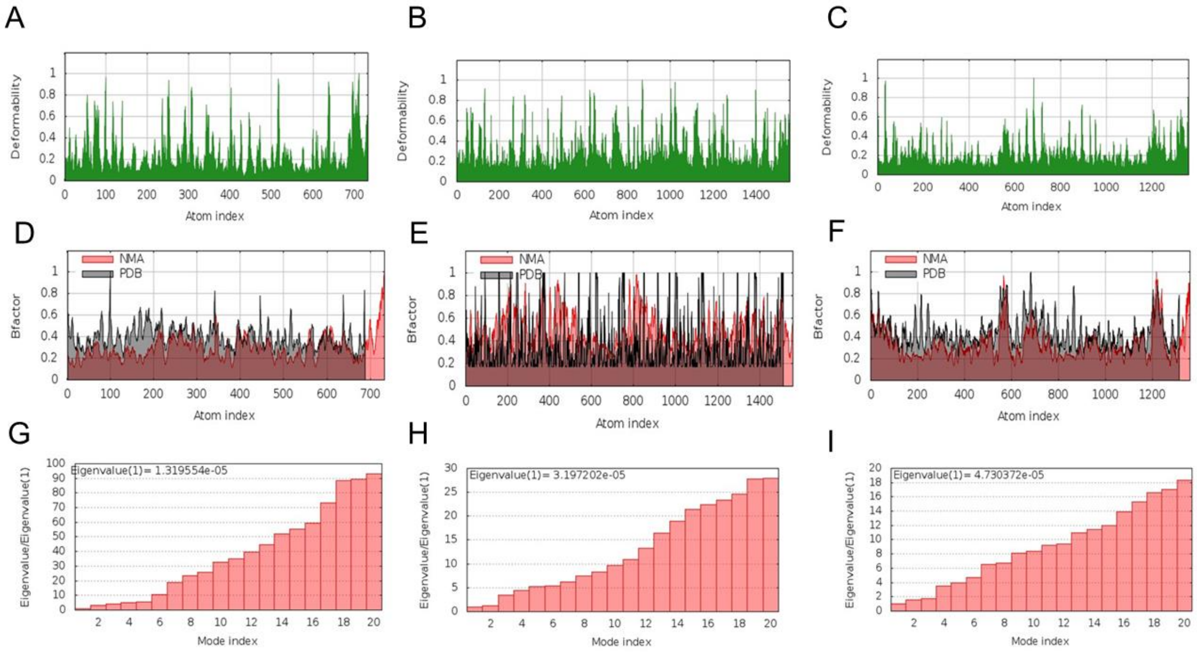


**Supplementary Fig. 3** Predicted conformational dynamics from iMODS. Deformability plot for peptide complex with **a** CTSD **b** BACE-2 **c** RNR. B-factor comparison for peptide complex with targets **d** CSTD **e** BACE-2 **f** RNR. Eigen-values for docked complex of peptide with **g** CSTD **h** BACE-2 **i** RNR.
